# Diverse microbial exposure exacerbates the development of allergic airway inflammation in adult mice

**DOI:** 10.1101/2025.03.21.644556

**Authors:** Jessica Elmore, Julie Sahler, Sabrina Solouki, Nicholas Koylass, Albert Wang, Sophie Nelissen, Amie Redko, Weishan Huang, Avery August

**Affiliations:** Cornell Center of Immunology, Cornell Institute for Host Microbe-Interactions and Disease, Department of Microbiology & Immunology, Cornell University College of Veterinary Medicine, Ithaca, NY, USA; Department of Pathobiological Sciences, Louisiana State University, Baton Rouge, LA, USA; Department of Biomedical Sciences, Cornell University, College of Veterinary Medicine, Ithaca, NY, USA; Cornell Center for Health Equity, Cornell University, Ithaca, NY, USA

**Keywords:** allergic asthma, hygiene hypothesis, pet store mice

## Abstract

**Background:** Exposure to a diversity of microbes has been implicated in playing a major role in susceptibility to the development of allergic lung type diseases. The hygiene hypothesis suggests that those exposed to a broad diversity of microbes are more likely to be protected against developing allergic type diseases. However, changes in exposure to microbial diversity can occur in both younger individuals, as well as in adults, and the effects are not always understood.

**Objective:** We investigated the effect of exposure to broad microbial diversity on the airway T cell response in house dust mite (HDM) induced allergic airway disease (AAD, a model of allergic asthma).

**Methods:** We increased exposure to broad microbial diversity by co-housing specific pathogen free (SPF) adult or newborn mice with pet store mice (PSE or BiPSE, respectively). Mice were then exposed to HDM to induce AAD.

**Results:** We found that the effect of increased microbial exposure on the development of allergic airway inflammation differs by age. Increasing exposure to diverse microbes as adults exacerbates the development of allergic airway inflammation, whereas this was not observed when exposure occurred at birth.

**Conclusion:** We suggest that experimental evaluation of the hygiene hypothesis in inflammation, particularly those using mouse models, may need to consider age of the host and time of microbial exposure.

**Capsule Summary:** Mouse models of increased exposure to diverse microbial environment shown to differentially affect the development of allergic airway inflammation, depending on the age of microbial exposure.

## Introduction

Lung inflammatory disorders such as allergic asthma are generally caused by hypersensitive reactions to antigens (1, 2). Allergic asthma can be triggered by environmental and genetic factors driven by type 2 innate lymphoid cells (ILC2s), T_H_2 cells, T_H_17 cells, neutrophils, or eosinophil responses (3). Sensitization to an allergen, such as house dust mite (HDM), followed by repeated airway challenge promotes type 2 asthma characterized by T_H_2 cytokines IL4, IL5, or IL13 (4). IL5 and IL13 production from T_H_2 and ILC2s promote lung eosinophilia, and IL13 can cause increases in mucus production and T_H_2 cell recruitment into the lungs (5-7). T_H_17 responses can also develop, and production of IL17A promotes neutrophil trafficking into the airways and can promote airway restriction (8). Such T_H_17/neutrophilic responses are associated with severe form of allergic asthma and can be resistant to corticosteroids (8, 9).

The effect of the microbiota on the host-immune response on lung inflammatory diseases and allergic asthma have been the focus of a number of studies (10-13). Previous work has examined the effects of shifts in the gut microbiota on mouse models of Allergic Airway Disease (AAD). Long term exposure to the narrow-spectrum antibiotic vancomycin, or streptomycin both prior to and after birth, led to increased susceptibility to developing allergic asthma in a model of ovalbumin-induced allergic airway disease (14, 15). Germ free mice, or those with limited microbial burden (specific pathogen free, SPF) are more susceptible to developing HDM-induced allergic asthma (15). In addition, differences in housing facilities and their environmental quality affected the severity of HDM induced airway disease, where mice in SPF facilities had increased levels of inflammation (13).

The leading theory for why increasing prevalence of allergies and allergic asthma is the hygiene hypothesis, first proposed in the 1980s (16). This hypothesis states that those who were exposed to “dirty” environments at an early age are protected against developing allergic type diseases in adulthood (17). The hygiene hypothesis has directed attention to the effects of the microbiome on different diseases (12), e.g., early childhood exposure to microbes correlates with a reduction in allergic and autoimmune diseases in later life (18, 19). A number of studies have shown that the gut microbiota affects the immune response, including in allergic diseases in mammalian hosts (11, 12, 20). Lifestyle changes such as weight fluctuations, dieting, obesity, or living environment(s) can all contribute to shifts or reduction in the diversity of microbial communities which can affect the way our immune system responds to disease (12, 21, 22).

SPF mice are the most commonly used models in laboratory research, however they have limited microbial burden that may affect their immune system and their responses to disease (23). Notably, SPF mice have been reported to lack tissue resident antigen specific memory T cells, or terminally differentiated effector memory cells, characteristics that are more associated with neonatal than adult human immune systems (24). Understanding the effect of the exposure to a variety of microbial antigens on the immune system may lead to a better understanding of disease models, which could further lead to better treatment options. Here we explore the consequences on lung inflammation in models of allergic asthma upon shifting the normal microbiota through exposure at different ages.

## Materials and Methods

### Mice

Wild type non-reporter (WT), or dual reporter IL17A-GFP/Foxp3-RFP, or IL10-GFP/Foxp3-RFP mice all on a C57Bl/6 background (25) were bred and maintained at Cornell University. Pet store mice were purchased from local pet stores in the Finger Lakes Region in New York. Mice were housed in SPF environment or cohoused with pet store mice in non-SPF environment. Mice between 6-12 weeks of age were used, and all experiments were approved by the Office of Research Protection’s Institutional Animal Care and Use Committee at Cornell University.

### Induction of HDM induced AAD

WT, IL17A-GFP/Foxp3-RFP mice were exposed to 10μg or 15μg HDM extract (from *D. pteronyssinus*; obtained from Greer Labs, item no. XPB70D3A2.5) intranasally as previously described for up to 10 consecutive days (26). Twenty-four hours after the last exposure, mice were euthanized and cell populations were analyzed via flow cytometry. IgE ELISA. Serum was collected IgE levels analyzed by enzyme-linked immunosorbent assay using IgE enzyme-linked immunosorbent assay kit (BD Biosciences) as previously describe (27).

### Enhancing exposure to broad microbial diversity by co-housing with pet store mice

SPF WT, IL17A-GFP/Foxp3-RFP mice or breeding pairs were transferred to a non-SPF facility and co-housed with pet store mice for up to 3 months. Bi-weekly bleeds were taken to monitor the emergence of T cell memory populations that resemble the adult human immune system as has been reported (24). These mice are given the designation of “converted” pet store exposed (PSE) after evaluation of increased T cell memory population compared to SPF mice. Additionally, pups born from these breeder pairs were continuously co-housed with pet store mice throughout the duration of experimentation. Mice that were born under these conditions were given the designation of “born in” pet store exposed (BiPSE). Cage droppings from PSE mice were collected for analysis of specific pathogens.

### Lymphocyte Isolation

Lymphocytes were isolated and analyzed by flow cytometry from bronchial lavage fluid (BAL) and lungs. Cells were stained with the following antibodies at 1:200 in staining buffer: Fixable Viability Dye eFluor 506 (Invitrogen; 65-0866-14), anti-Ly6G PE-Cy7 (RRID:AB_2811793) or eFluor 450 (RRID:AB_2637124), anti-CD117 FITC (RRID:AB_465186), anti-SiglecF PE (RRID:AB_2637129), anti-CD11b PE-Texas Red (RRID:AB_10373548), anti-CD11c APC (RRID:AB_117310), anti-MHC II AF700 (RRID:AB_494009), anti-CD49b PerCP eF710 (RRID:AB_11150237), anti-FcεRIα PECy7 (RRID:AB_10640122), anti-F4/80 APC/Cy7 (RRID:AB_893477), anti-IL17A PerCP-CY5.5 (RRID:AB_925753) or FITC (RRID:AB_2534391) (eBioscience), anti-TCRβ APC-CY7 (Biolegend; RRID:AB_893624), anti-CD4 Alexa Fluor 700 (RRID:AB_493999) or eFluor 450 (RRID:AB_10718983), anti-CD8α PE-Texas Red (RRID:AB_11152075) or PerCP-Cy5.5 (RRID:AB_2075238, anti-TCRγδ APC (RRID:AB_1731824), anti-NK1.1 Allophycocyanin (RRID:AB_389363), anti-CD44 V500 (RRID:AB_1937328), anti-CD62L PE-Cy7 (RRID:AB_313103), anti-CD11b V450 (BD Biosciences; RRID:AB_1645266) or PE-Dazzle594 (Biolegend; RRID:AB_2563648) anti-B220 Alexa Fluor 700 (Biolegend; RRID:AB_493717), anti-CD8 (BD Biosciences; RRID:AB_2732919) or CD4 BUV395 (BD Biosciences; RRID:AB_2738426), Fc blocking antibody was from eBioscience. Intracellular cytokine production was determined using PMA (Sigma-Aldrich P1585-1MG) and Ionomycin (Sigma-Aldrich I0634-1MG) stimulation. Cells were analyzed using BD FACS Aria II or BD FACSymphony and analyzed with FlowJo software.

### Histology

Lung sections were stained with H&E or PAS and imaged with a 20X objective on the Zeiss AX10 imager.m1 microscope and camera system.

### Statistical analysis

For data analysis, one-way ANOVA or Mann Whitney test were performed using GraphPad Prism version 8.00 or 9.01 (GraphPad, San Diego, CA). Statistically significant differences have the probability of *p* ≤ 0.05.

## Results

### Exposure to broad microbial diversity as adults is associated with enhanced allergic airway induced inflammation

In order to examine a mouse model of exposure to broad diversity of microbial flora, adult mice were housed with pet store mice that carry a wide range of microbial organisms to generate pet store exposed (PSE) mice (see **Supplemental Table 1** for example of microbial burden) (24). Weights of the PSE mice were monitored and blood was collected weekly for analysis of immune cell changes. PSE mice exhibited reduced survival over the 80d exposure time (**Supplemental Fig. 1A**), and lost a significant amount of weight within the first 10d of co-housing (**Supplemental Fig. 1B**). This suggests that as previously reported, cohousing of SPF mice with pet store mice led to increased microbial and pathogen burden over this time (24). Similarly, as previously reported, the overall population of CD8^+^ central (CD62L^+^CD44^+^ CM) and effector memory (CD62L^-^CD44^+^ EM) T cells was higher than the SPF mice (time 0) after 2 weeks of exposure to pet store mice, and was maintained through 65 days of cohousing (**Supplemental Fig. 1C-F**). These results indicate that the immune systems of the PSE mice have “converted” to resemble an “adult human” immune system phenotype as has been previously reported for this model (24, 28, 29). We used this mouse model of broad microbial exposure to examine whether this different immune state affected the pathogenesis of lung inflammation in AAD. Adult SPF cytokine reporter mice (IL17A-GFP/Foxp3-RFP and IL10-GFP/Foxp3-RFP mice) either remained in SPF facilities as controls, or were transferred to non-SPF facilities and co-housed with pet store mice. In addition, since it has been reported that exposure to a variety of microbes early in life protects against developing allergic diseases later in life (30), we also wanted to compare to mice exposed to broad diversity of microbial flora prior to and after birth. To that end, breeder pairs of SPF IL17-GFP/Foxp3-RFP and IL10-GFP/Foxp3-RFP mice were co-housed with pet store mice, and their pups (referred to as born in PSE mice (BiPSE)) were also continuously co-housed with pet store as they aged.

SPF or PSE mice were exposed to HDM to induce allergic airway induced inflammation, or PBS as control, and lung cell populations were analyzed via flow cytometry. We found that PSE mice exposed to HDM to induce AAD exhibited increased lung inflammation as determined by lung histology compared to HDM exposed SPF mice (**Fig. 1A**). Exposure of SPF mice to HDM led to an increase in overall serum IgE as expected, however, PSE mice exhibited elevated serum IgE in the absence of HDM exposure, suggesting that exposure to broad microbial diversity increased basal IgE, which was not further increased with exposure to HDM (**Supplemental Fig. 2**). Total lung cell numbers, and total lung T cell numbers were significantly increased in HDM exposed PSE mice compared to HDM exposed SPF mice (**Fig. 1B**). Among the CD4^+^ T cells, the percentage of T_H_17 cells (determined via IL-17A/GFP expression) was significantly increased in HDM exposed PSE mice compared to HDM exposed SPF mice (**Fig. 1C**). While HDM exposure led to an increase in the percentage of Foxp3^+^ T regulatory cells (determined via Foxp3/RFP expression) in both SPF and PSE mice, there was no significant difference in the percentages between the two conditions (**Fig. 1C**). By contrast, HDM exposure also led to an increase in the percentage of Foxp3^-^ Type 1 regulatory cells (Tr1 cells, determined via Foxp3/RFP^-^ and IL10/GFP^+^ expression) in both SPF and PSE mice, but there was a reduced percentage of Tr1 cells in PSE mice, as well as reduced percentage of IL10^+^ Foxp3^+^ T regulatory cells (determined via Foxp3/RFP^+^ and IL10/GFP^+^ expression) in PSE mice (**Fig. 1C**).

**Figure 1.**
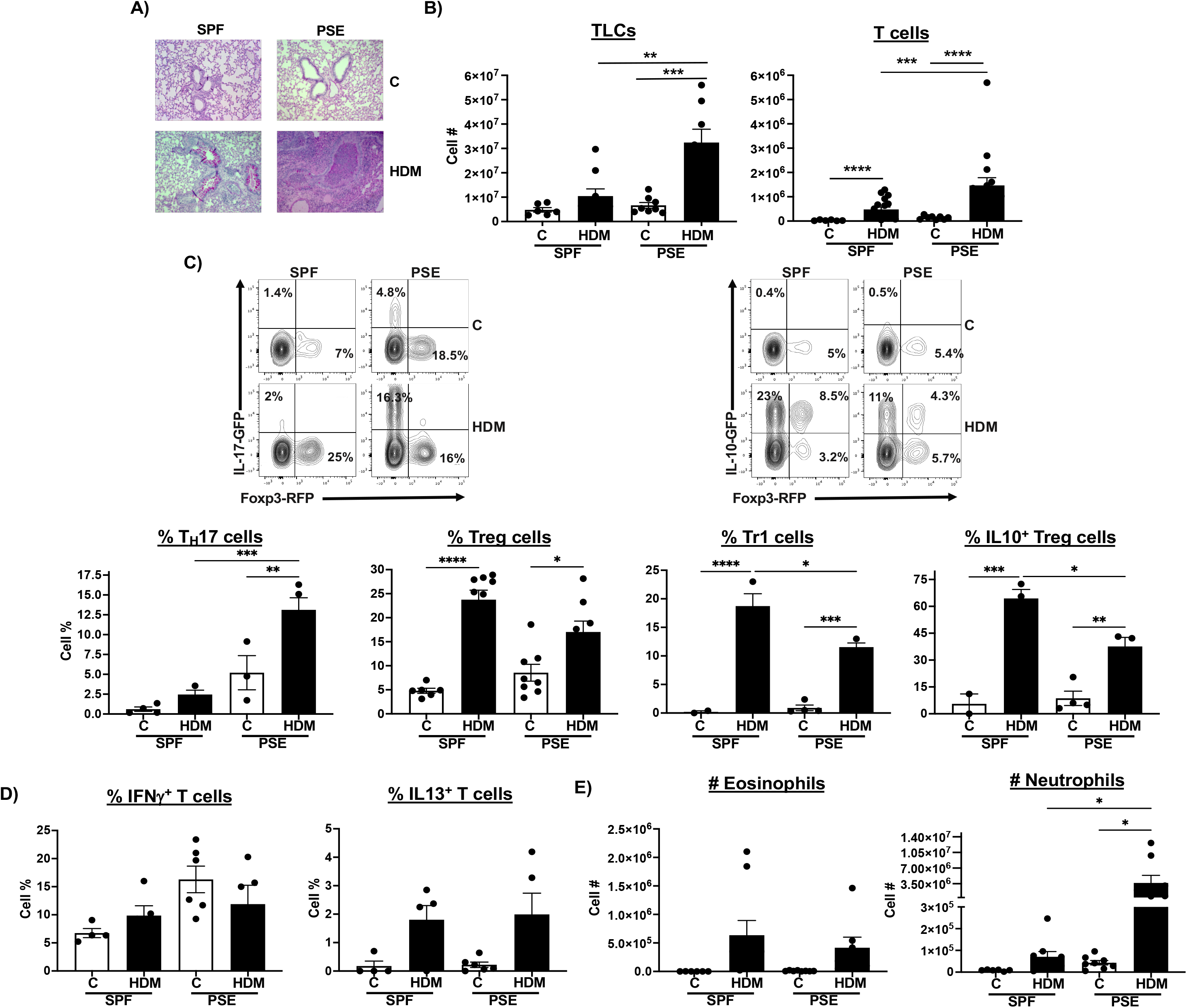
Enhanced inflammation in HDM induced AAD following exposure to broad diversity of microbes. SPF or PSE mice were exposed to PBS (C) or HDM for 10d. (**A**) Lung section tissues from PBS or HDM exposed SPF or PSE mice were stained with PAS. **(B)** Numbers of total lung cells (left: SPF/C: n=3; SPF/HDM: n=9; PSE/C: n=8; PSE/HDM: n=8) or total lung T cells (right: SPF/C: n=6; SPF/HDM: n=18; PSE/C: n=8; PSE/HDM: n=16, **p<0.01, ***p<0.001, ****p<0.0001, Mann-Whitney test) from control or HDM exposed mice. (**C**) Top from left to right: example flow cytometric plots of analysis of lung CD4^+^ T cells to identify T_H_17 (IL17A-GFP^+^/Foxp3-RFP^-^) or Foxp3^+^ Tregs (IL17A-GFP^-^/Foxp3-RFP^+^) (left), or Tr1 (IL10-GFP^+^/Foxp3-RFP^-^), IL10^+^ Foxp3^+^ Tregs (IL10-GFP^+^/Foxp3-RFP^+^)(right). Bottom from left to right: percent lung T_H_17 (SPF/C: n=4; SPF/HDM: n=3; PSE/C: n=3; PSE/HDM: n=4), Foxp3^+^ Tregs (SPF/C: n=6; SPF/HDM: n=9; PSE/C: n=8; PSE/HDM: n=9), Tr1 (SPF/C: n=2; SPF/HDM: n=3; PSE/C: n=4; PSE/HDM: n=3), and IL10^+^ Foxp3^+^ Tregs (SPF/C: n=2; SPF/HDM: n=3; PSE/C: n=4; PSE/HDM: n=3), *p<0.05, **p<0.01, ***p<0.001, ****p<0.0001, one-way ANOVAs with Tukey’s multiple corrections test. **(D)** IFNγ (left) and IL13 production (right) production by lung CD4^+^ T cells was determined by PMA/Ionomycin stimulation *in vitro* and flow cytometric analysis. SPF/C: n=4; SPF/HDM: n=5; PSE/C: n=6; PSE/HDM: n=5. **(E)** Number of lung eosinophils (left) and neutrophils (right), SPF/C: n=6; SPF/HDM: n=9; PSE/C: n=8; PSE/HDM: n=8. *p<0.05. Ordinary one-way ANOVAs with Tukey’s multiple corrections test, data +/-SEM.

Neither SPF nor PSE mice exhibited a significant increase in IFNγ producing CD4^+^ T cells upon HDM exposure, and both had increases in IL13 producing CD4^+^ T cells (all determined via PMA/Ionomycin stimulation *in vitro* and intracellular cytokine staining) (**Fig. 1D**). Furthermore, the number of lung neutrophils was increased in HDM exposed SPF mice, and significantly increased in PSE mice compared to HDM exposed SPF mice (**Fig. 1E**). There was no difference in the HDM induced increase in the number of eosinophils in SPF and PSE mice (**Fig. 1E)**. These findings suggest that an effect of the increased microbial exposure is the development of T_H_17/neutrophil rich AAD. Taken together, these data suggest that exposure to diverse microbes led to enhanced HDM induced lung inflammation in AAD. These findings also suggest that increased microbial exposure may not always confer protection against developing allergic lung diseases, as predicted by the hygiene hypothesis (16).

### PSE mice induced to develop AAD are less sensitive to dexamethasone treatment

Allergic asthma is conventionally treated with corticosteroids, among other agents (31), however severe forms of asthma can be associated with T_H_17 cells, which are associated with steroid resistance (9, 32). Our data suggest that HDM induced AAD results in enhanced inflammation in the airways and increased T_H_17 cells in PSE mice to compared to SPF mice (see **Fig. 1**). We therefore wanted to determine whether the inflammation associated with elevated T_H_17 cells in PSE mice is susceptible to steroid treatment. To examine this, SPF and PSE IL17A-GFP/Foxp3-RFP mice were injected intraperitoneally with 1 mg/kg of dexamethasone (Dex) or vehicle control prior to the start of, and each day during 10 day HDM exposure (33). Lungs were collected and sections analyzed for lung inflammation and mucous production, which suggests that dexamethasone treatment reduced mucus production and pathology in both SPF and PSE mice (**Fig. 2A**).

**Figure 2.**
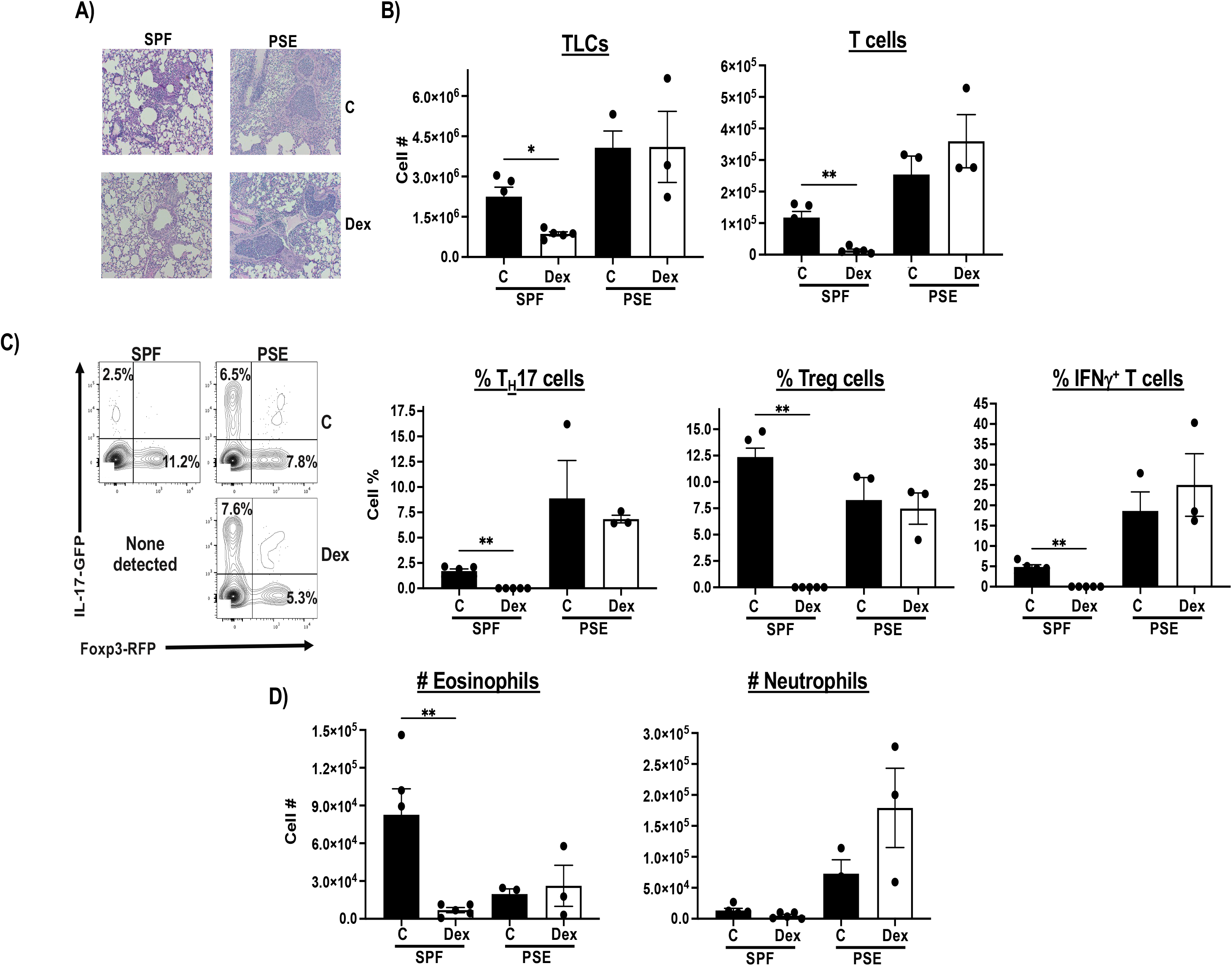
Effect of Dexamethasone treatment on HDM-induced lung inflammation in PSE mice. SPF or PSE mice were exposed intranasally to 10 μg HDM for 10 consecutive days, along with treatment (Dex) or vehicle control (C). (**A**) Top: Lung tissue sections were stained with PAS to detect inflammation and mucous. (**B**) Total lung cells (***left***), and total lung T cells (***right***). (*p<0.05, **p<0.01, Mann Whitney test). (**C**) Left: example flow cytometric plots of analysis of lung CD4^+^ T cells to identify T_H_17 (IL17A-GFP^+^/Foxp3-RFP^-^) and Foxp3^+^ Tregs (IL17A-GFP^-^/Foxp3-RFP^+^). ***Bottom left***: percent lung T_H_17, ***Bottom middle***: and Foxp3^+^ Tregs. ***Bottom right***: Percent IFNγ positive lung CD4^+^ T cells as determined by PMA/Ionomycin stimulation *in vitro* and flow cytometric analysis. **p<0.01, Mann Whitney test. (**D**) Number of lung eosinophils (left) and neutrophils (right), SPF/HDM/C: n=5; SPF/HDM/Dex: n=5; PSE/HDM/C: n=3; PSE/HDM/Dex: n=3, **p<0.01, Mann Whitney test, data +/-SEM.

Total lung cell numbers as well as total lung T cell numbers were significantly reduced by Dex treatment in SPF mice, but not in PSE mice, suggesting some steroid resistance (**Fig. 2C**). Correspondingly, the percentage of T_H_17 cells (determined via IL-17A/GFP expression) was significantly reduced by Dex treatment of HDM exposed SPF mice, but not in HDM exposed PSE mice (**Fig. 2B**). Similarly, Dex treatment of HDM exposed SPF mice led to a significant reduction in the percentage of Foxp3^+^ T regulatory cells (determined via Foxp3/RFP expression), as well as IFNγ producing CD4^+^ T cells (determined via PMA/Ionomycin stimulation *in vitro* and intracellular cytokine staining), but not in HDM exposed PSE mice (**Fig. 2C**). Furthermore, Dex treatment did not affect the number of lung neutrophils, nor eosinophils, in HDM exposed PSE mice (**Fig. 2D**). Overall, these findings suggest that PSE mice develop AAD that exhibits some resistance to corticosteroid treatment.

### Exposure to broad microbial diversity as newborns is associated with reduced allergic airway induced inflammation

We next evaluated mice exposed to broad diversity of microbial flora prior to birth. To that end, breeder pairs of SPF IL17-GFP/Foxp3-RFP and IL10-GFP/Foxp3-RFP mice were co-housed with pet store mice, and their pups (referred to as born in PSE mice (BiPSE)) were also continuously co-housed with pet store as they aged. At 6-8 weeks old, SPF or BiPSE mice were exposed to HDM to induce AAD, or PBS as control, and lung cell populations were analyzed via flow cytometry. We found that BiPSE mice exposed to HDM to induce AAD exhibited similar lung inflammation as determined by lung histology compared to HDM exposed SPF mice (**Fig. 3A**). Furthermore, unlike what was observed in HDM exposed PSE mice, there was no significant difference in the number of total lung cells, or in numbers of total T cells between HDM exposed SPF and BiPSE mice (**Fig. 3B**). The percentage of CD4^+^ T_H_17 cells (determined via PMA/Ionomycin stimulation *in vitro* and intracellular cytokine staining) was increased in both HDM exposed SPF and BiPSE mice, but again, unlike what was observed for HDM exposed PSE mice, there was no difference between HDM exposed SPF and BiPSE mice (**Fig. 3C**). Similarly, HDM exposure led to an increase in the percentage of Foxp3^+^ T regulatory cells (determined via Foxp3/RFP expression) in both SPF and BiPSE mice, there was no significant difference in the percentages between the two conditions (**Fig. 3C**). In addition, HDM exposure induced an increase in the percentage of Tr1 cells (determined via Foxp3/RFP^-^ and IL10/GFP^+^ expression) in both SPF and BiPSE mice, as well as reduced percentage of IL10^+^ Foxp3^+^ T regulatory cells (determined via Foxp3/RFP^+^ and IL10/GFP^+^ expression), unlike what was observed for PSE mice, there was no difference in this induction between SPF and BiPSE mice (**Fig. 3C**). Furthermore, neither SPF nor BiPSE mice exhibited a significant difference in IFNγ producing CD4^+^ T cells upon HDM exposure (determined via PMA/Ionomycin stimulation *in vitro* and intracellular cytokine staining) (**Fig. 3D**). Notably, BiPSE mice had significantly higher number of neutrophils at baseline compared to SPF mice (p=0.032 by t test), which was not further increased after HDM exposure mice (**Fig. 3E**). Both SPF and BiPSE mice developed HDM induced increase in the number of eosinophils (**Fig. 3E)**. These findings suggest that unlike the case for exposure to increased microbial exposure as adults, increased microbial exposure at birth does not lead to enhanced AAD, but also did not confer protection as predicted by the hygiene hypothesis (16)(34).

**Figure 3.**
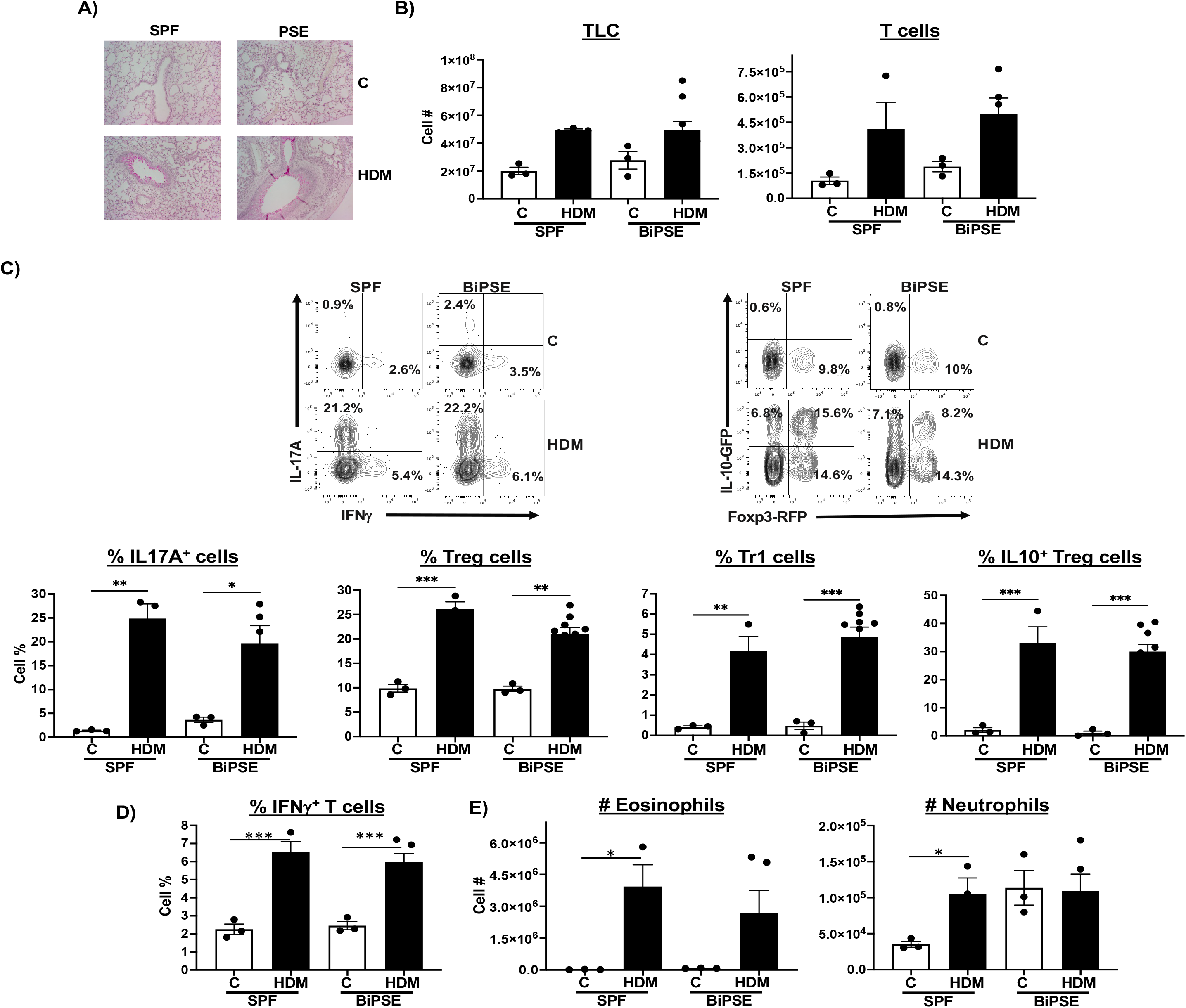
BiPSE mice do not exhibit enhanced inflammation in HDM induced AAD following exposure to broad diversity of microbes. SPF or BiPSE mice were exposed to PBS or HDM for 10d. (**A**) Lung section tissues from PBS or HDM exposed SPF or BiPSE mice were stained with PAS. **(B)** Numbers of total lung cells (left) or total lung T cells (right) from PBS control (C) or HDM exposed mice. For total lung cells: SPF/C: n=3; SPF/HDM: n=3; BiPSE/C: n=3; BiPSE/HDM: n=9. For total lung T cells: SPF/C: n=3; SPF/HDM: n=3; BiPSE /C: n=3; BiPSE /HDM: n=5. (**C**) Top: Example flow cytometric plots of analysis of lung CD4^+^ T cells to identify: T_H_17 (IL17A-GFP^+^/Foxp3-RFP^-^) or Foxp3^+^ Tregs (IL17A-GFP^-^/Foxp3-RFP^+^) (left), or Tr1 (IL10-GFP^+^/Foxp3-RFP^-^), IL10^+^ Foxp3^+^ Tregs (IL10-GFP^+^/Foxp3-RFP^+^), IL10^-^ Foxp3^+^ Tregs (IL10-GFP^+^/Foxp3-RFP^+^), or total Foxp3^+^ Tregs (IL10-GFP^+/-^/Foxp3-RFP^+^) (right). Bottom from left to right: percent lung T_H_17 (SPF/C: n=3; SPF/HDM: n=3; BiPSE/C: n=3; BiPSE/HDM: n=5), Foxp3^+^ Tregs, Tr1, and IL10^+^ Foxp3^+^ Tregs (SPF/C: n=3; SPF/HDM: n=3; BiPSE/C: n=3; BiPSE/HDM: n=9, *p<0.05, **p<0.01, ***p<0.001, one-way ANOVA). **(D)** IFNγ production by lung CD4^+^ T cells as determined by PMA/Ionomycin stimulation *in vitro* and flow cytometric analysis. SPF/C: n=3; SPF/HDM: n=3; BiPSE/C: n=3; BiPSE/HDM: n=5, ***p<0.001. **(E)** Number of lung eosinophils (left) and neutrophils (right), SPF/C: n=3; SPF/HDM: n=3; BiPSE/C: n=3; BiPSE/HDM: n=5, *p<0.05 by t test. data +/-SEM.

## Discussion

The hygiene hypothesis, first proposed in the 1980s, stated that exposure to “dirty” environments at a young age decreases susceptibility to developing allergic-type diseases in adulthood. A major attention of the hygiene hypothesis is the microbiome, and exposure of children to a broad diversity of microbes in their early life, leading to changes in the microbiota, exhibit effects on the development of their immune system (35). A number of studies have shown that the gut microbiota affects the immune response, including in allergic-type diseases in mammalian hosts (12, 14, 20). Germ-free mice have been shown to exhibit increased susceptibility to developing allergic asthma compared to SPF mice (36). However, while SPF mice carry a diverse microbiome compared to germ-free mice, they been shown to have immune systems that resemble that of newborns, which could greatly impact their immune responses to any inflammatory challenge. Exposing them to a broader diversity of microbiota lead to changes of their immune system to more resemble adult humans (24), and alters their immune responses. Here we explored the consequences of exposure of mice to increased diversity of microflora on a model of allergic airway disease. Our findings suggest that the effect of exposure to broad microbial diversity may have an exacerbating effect, dependent on age of exposure to such diversity.

Reducing microbial burden through antibiotics is relatively easy and a well-controlled means to decrease microbiome diversity, however the reverse is more difficult to modify and control. Cohousing of SPF mice with pet store mice adds more realistic exposures, but also more variability, such as 1) microbiomes of the pet store mice may not be identical between pet stores or even the same pet store purchased over time, 2) cohousing of different mice occasionally leads to fighting and exterior wounds that could result in unintended inflammation, 3) mental stress of new social dynamics could influence hormones or other cell functions, and 4) determining the duration of cohousing before beginning an experiment is still debated. We have circumvented some of these variables by excluding mice with obvious wounds, waiting a minimum of 5 weeks post cohousing for social structures to balance out, and testing for conversion to determine immune memory cell percentages in blood to establish mouse eligibility for experiments.

Our results indicated that increasing exposure to microbial diversity in adult SPF mice (PSE mice) resulted in enhanced development of a T_H_17/neutrophil rich lung inflammation, analogous to T_H_17/neutrophilic type asthma, associated with a more severe form of asthma (37). T_H_17 cell responses in asthma are associated with corticosteroid resistance, a treatment often used for moderate to severe asthma patients (38, 39). Treatment of HDM-induced AAD PSE mice with Dexamethasone revealed that these mice with T_H_17/neutrophilic type airway inflammation were less sensitive to the corticosteroid than SPF mice. Overall, increased diversity in microbiota in this model skewed their inflammation towards an T_H_17/neutrophilic type AAD response that was less sensitive to corticosteroids.

By contrast, mice that were born in the presence of pet store mice (BiPSE mice) such that their exposure to broad microbial diversity occurred at birth exhibited no difference in the development of AAD compared to SPF mice. This suggests that mice born in the presence of increased microbial diversity (conferred by co-housing with the pet store mice) have altered inflammatory responses compared to adult PSE mice when they are exposed to HDM. It is likely that exposure to broad microbial diversity as adults shape the immune system of the mice such that they respond with an enhanced T_H_17/neutrophilic type AAD. Mice exposed to broad microbial diversity as newborns may have mechanisms, such as induction of tolerance, that may shape their immune systems in a different manner, and thus drive a different response to HDM exposure. Our findings that mice exposed to conditions of broader microbial diversity exhibit elevated AAD is contrary to what has been proposed by the hygiene hypothesis. Instead, specific microbiome composition may have a larger influence than simply total microbial diversity. Generating microbiome signatures, and then associating those with types of inflammation will be crucial to understand the links between the two.

Although we did not focus on in-depth analysis of specific microbial communities, we found that increasing microbial diversity may alter allergic lung disease outcomes. Overall, our findings suggest that the age at which mice are exposed to broad microbial diversity may determine their response in lung inflammatory diseases.

## Abbreviations

AAD: Allergic Airway Disease
BiPSE: born-in pet store exposed
CM: central memory
Dex: dexamethasone
EM: effector memory
HDM: house dust mite
ILCs: Innate Lymphoid cells
PBS: Phosphate-Buffered Saline
PSE: pet store exposure
T_H_17: T helper 17 cells
TLC: Total lung lymphocytes
TSLP: Thymic Stromal Lymphopoietin
SPF: specific pathogen free

## Acknowledgements

We would like to thank members of the August lab for their feedback and the Cornell Center for Animal Resources and Education for housing pet store mice.

## Figure legends

**Supplemental Figure 1. Exposure of SPF mice to pet store mice generates a more mature CD8**^**+**^ **T cell memory population**. SPF mice were co-housed with pet store mice over 80 days. Mice were monitored for their (**A)** survival and (**B**) weight loss. n=13, *p<0.05, unpaired t test. Blood was collected each week to determine (**C-F**) CD8^+^ memory cell populations via flow cytometry gating of CD8^+^ memory cell populations with example gating shown in (**C**). CM: central memory; EM: effector memory; n=10 (day 0), n=2 (day 8), n=8 (day 16), n=5 (day 22), n=12 (day 36), n=7 (days 50 & 64), Data +/-SEM.

**Supplemental Figure 2. Exposure of SPF mice to pet store mice generates higher basal elevated serum IgE**. SPF or PSE mice were exposed to PBS (C) or HDM for 10d as in figure 1. (**A**) Serum IgE levels were determined. SPF/C: n=6; SPF/HDM: n=5; PSE/C: n=8; PSE/HDM: n=4. **p<0.01, ***p<0.001, Ordinary one-way ANOVAs with Tukey’s multiple corrections test. Data +/-SEM.

**Supplemental Table 1. Analysis of microbial agents in mouse cages**. Cage droppings from SPF or PSE mice were analyzed for the indicated intestinal microbial agents. n ≥ 6/group.

