## Supplementary figures and images for "Diverse microbial exposure exacerbates the development of allergic airway inflammation in adult mice"

### Supplemental figures

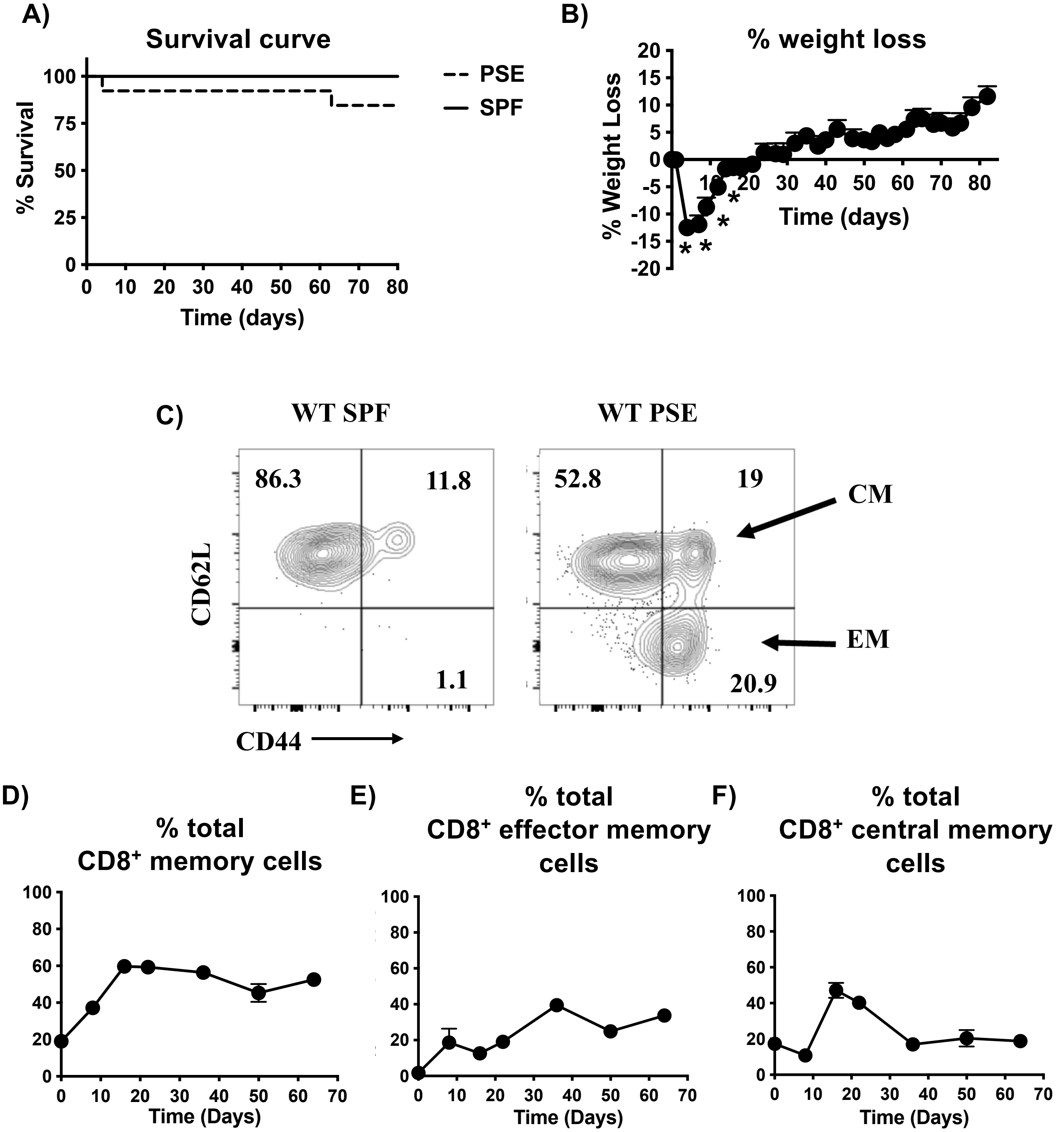

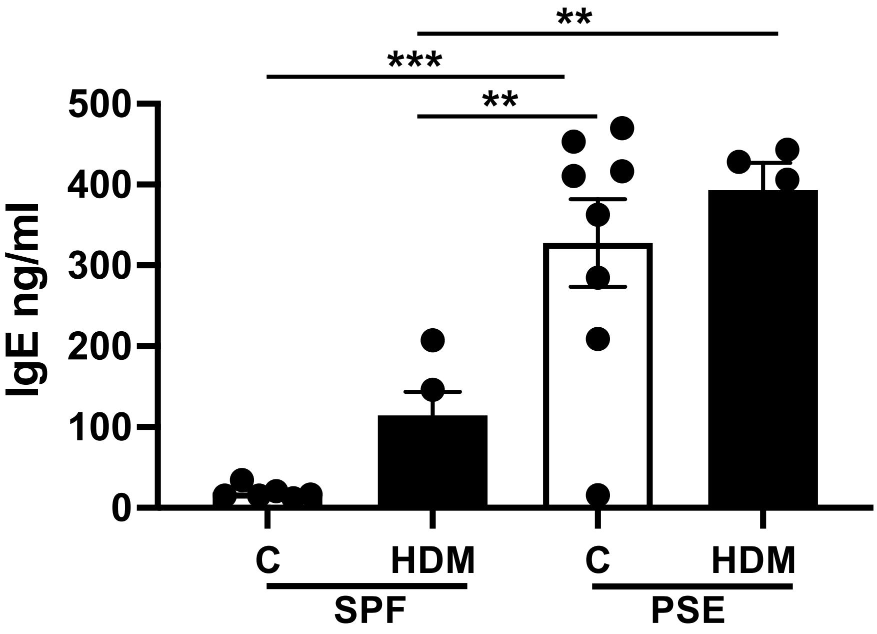
